## Supplementary Figure and Table for "HoxB-derived hoxba and hoxbb clusters are essential for the anterior-posterior positioning of zebrafish pectoral fins"

### Supplementary Fig. 1

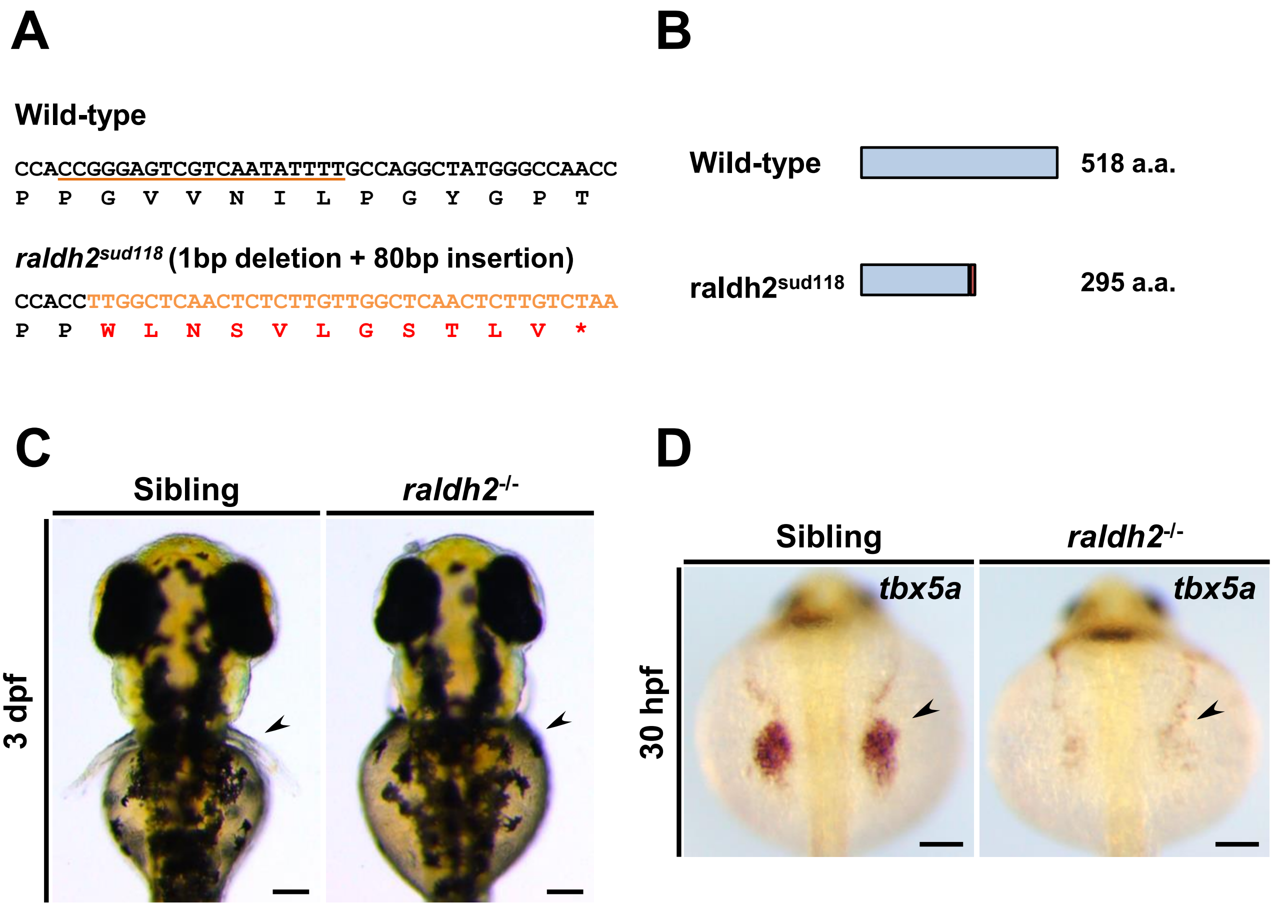


**Figure S1. Generation of the zebrafish *raldh2* mutants by CRISPR-Cas9**. (A) The nucleotide sequence surrounding the mutations in *raldh2* is shown on the left. The target sequence of the crRNA is emphasized with an underline, while the inserted nucleotide sequence is indicated in orange. Below the DNA sequence, the predicted amino acid is displayed, with red letters highlighting the abnormal amino acid resulting from the frameshift mutation. The asterisk represents the termination codon. (B) A schematic representation of the predicted protein structure is shown, with the red box indicating the abnormal amino acid sequences resulting from the frameshift mutations. The total number of predicted amino acids is displayed to the right of the schematic. (C) The absence of pectoral fins (arrowhead) in *raldh2* homozygous mutants is observed dorsally at 3 dpf. (D) The expression of *tbx5a* in the pectoral fin buds (arrowhead) is significantly reduced in *raldh2* mutants. The phenotype of our *raldh2*$^{sud118}$ mutants closely resembles the phenotype observed in previously isolated *raldh2* mutants.

### Supplementary Fig 2

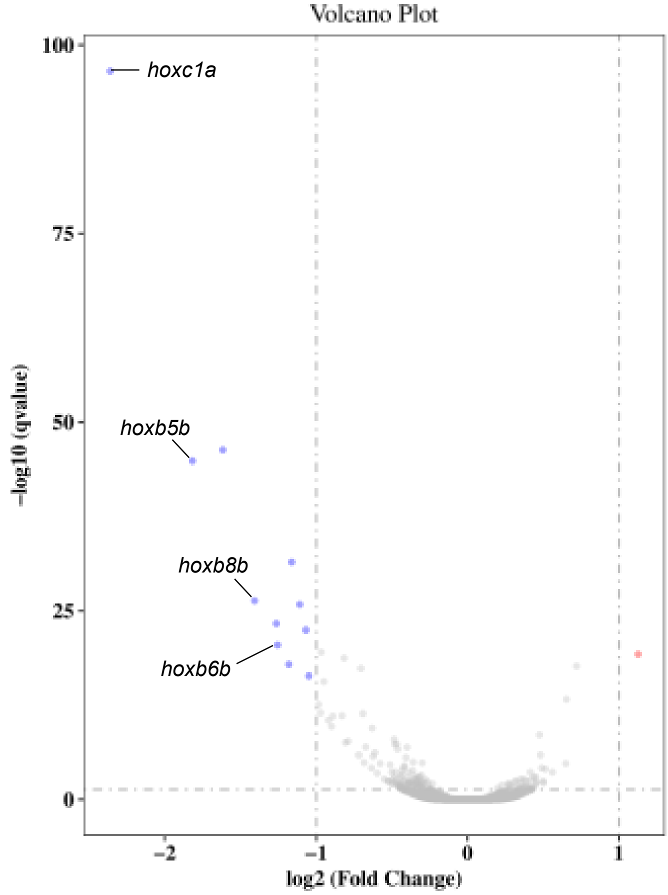


**Figure S2. Volcano plot of the transcriptome analysis between sibling and *raldh2* mutants**. The volcano plot illustrates both upregulated and downregulated differentially expressed genes from the comparison between wild-type and *raldh2* mutant embryos. Dot with more than a 2-fold increase in expression is indicated in orange, while those with more than a 2-fold decrease are shown in light blue. Among them, dots corresponding to *hox* genes are specifically highlighted.

### Supplementary Figure 3

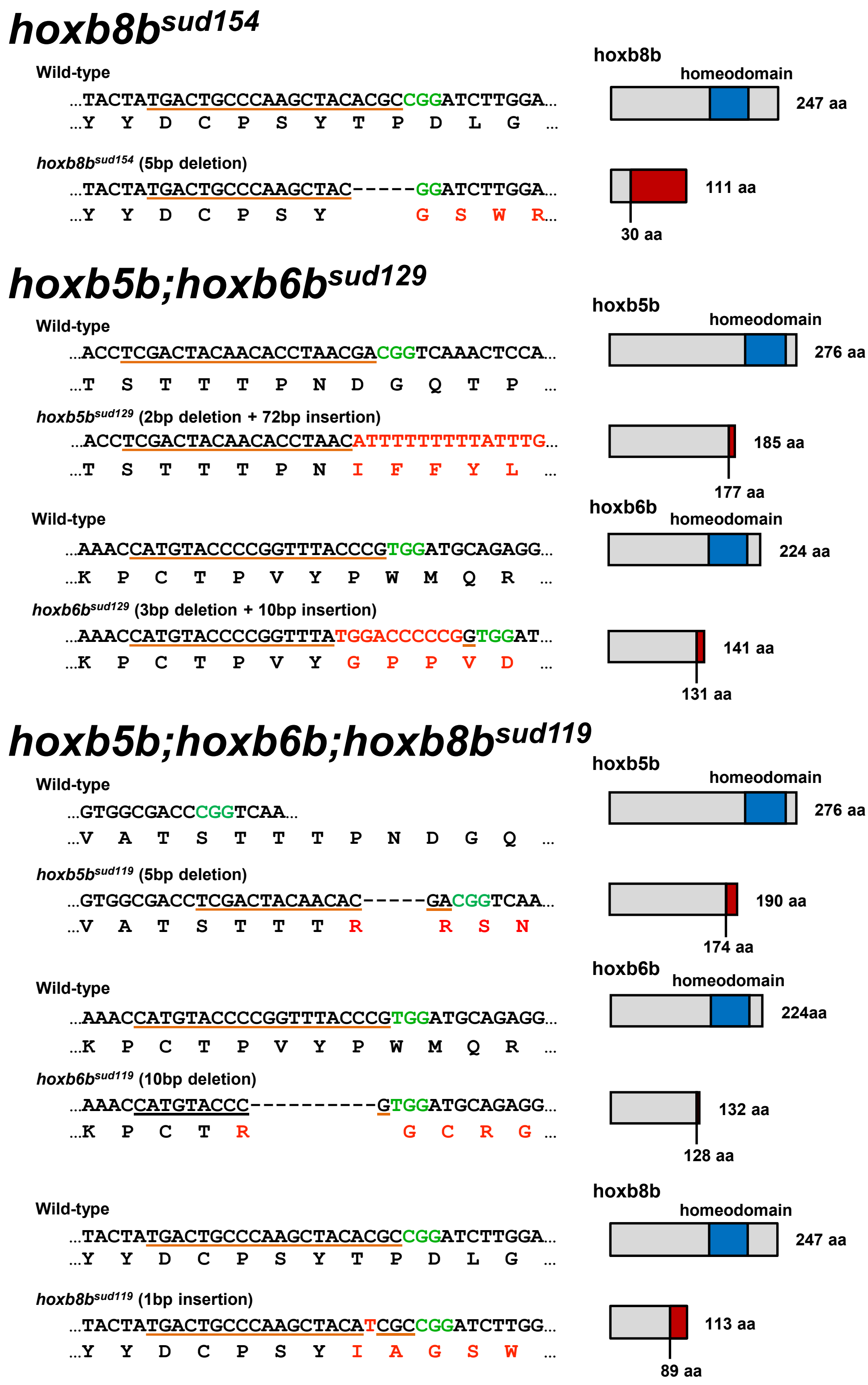


**Figure S3. Generation of the frameshift-induced hox mutants using CRISPR-Cas9**. Schematic representations show zebrafish *hoxb8b, hoxb5b;hoxb6b,* and *hoxb5b;hoxb6b;hoxb8b* mutants. The other frameshift-induced hox mutants were previously described (Maeno et al., 2024). The nucleotide sequence around the mutations is shown on the left. The target sequence of crRNA is emphasized with an orange underline. The inserted nucleotide sequence is emphasized in red. Below the DNA sequence, the predicted amino acid is shown. Red letters indicate the abnormal amino acid caused by the frameshift mutation. On the right, a schema represents the predicted protein structure. The blue box indicates the homeodomain, and red box indicates the abnormal amino acid sequences from the frameshift mutations. The total number of the predicted amino acids is shown on the right of the schema.

### Supplementary Figure 4

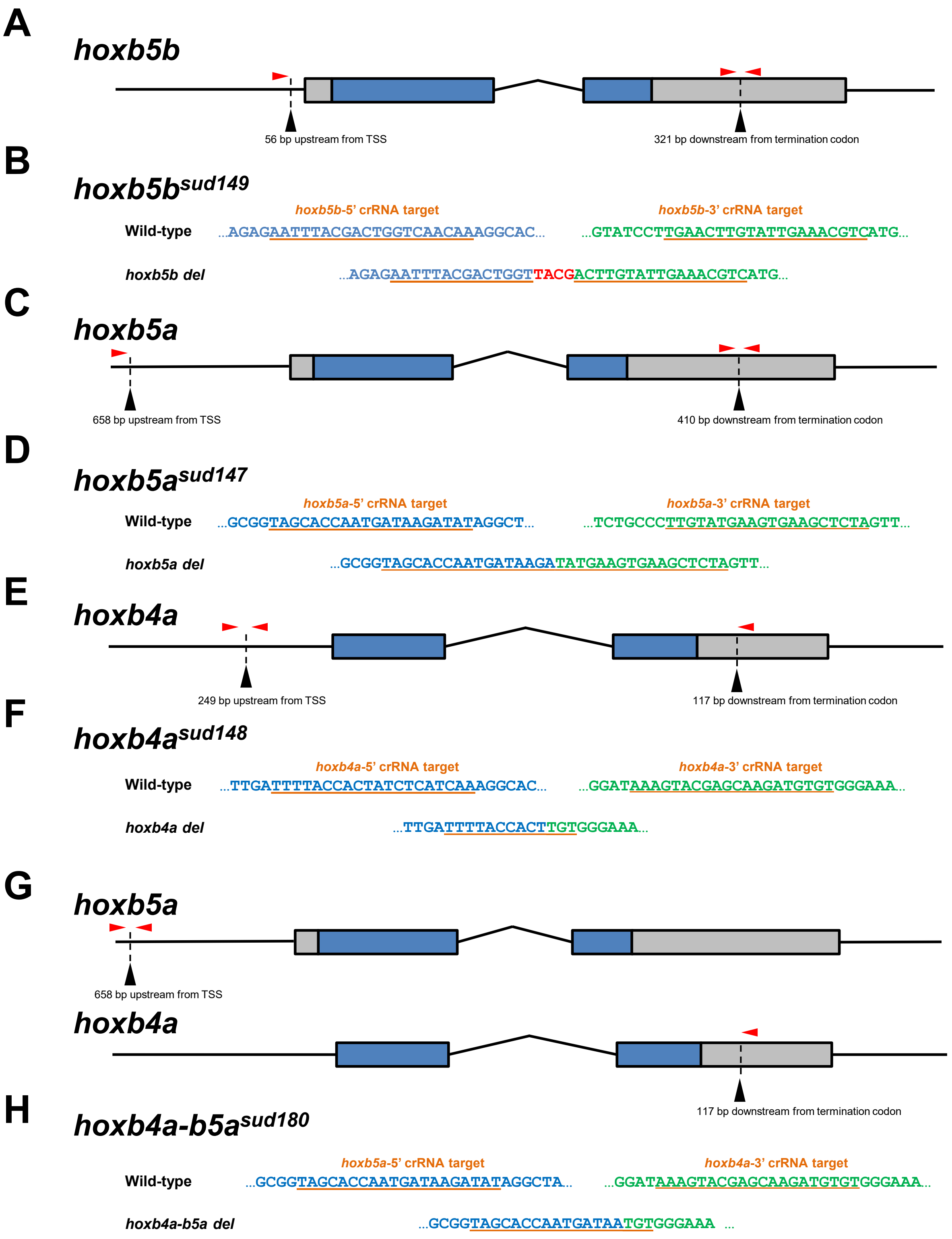


**Figure S4. Generation of locus-deletion mutants using CRISPR-Cas9**. (A, C, E, G). Schematic representations show the genomic structure of zebrafish *hoxb4a, hox5a,* and *hoxb5b*. Boxes represent the exons, with the blue boxes indicating the coding regions. Black arrowheads indicate the target sites of crRNA, while red arrowheads represent the primers used for genotyping. (B, D, F, H) Display genomic locus deletions of *hox4a, hox5a, hoxb5b,* and *hoxb4a-b5a* deletion mutants. The flanking sequences of the 5'- and 3'-crRNA targets (underline) are shown in blue and green letters, respectively.

#### Table S1 The frameshift mutations introduced in *hoxbb* genes in the embryos lacking pectoral fins during the screening

| No | Phenotype | Parents (male) | Parents (female) | *hoxba* | *hoxbb* | *hoxb5b* | *hoxb6b* | *hoxb8b* |
|---|---|---|---|---|---|---|---|---|
| 1 | No pectoral fin | *hoxba* $^{+/-}$ ;*hoxbb* $^{+/-}$ | *hoxba+/-;b5b, b6b, b8b* gRNA-injected | -/- | +/- | frameshift (2bp ins) | frameshift (1bp ins) | frameshift (5bp del) |
| 2 | No pectoral fin | *hoxba* $^{+/-}$ ;*hoxbb* $^{+/-}$ | *hoxba+/-;b5b, b6b, b8b* gRNA-injected | -/- | +/- | frameshift (5bp del) | frameshift (10bp del) | frameshift (1bp del) |
| 3 | No pectoral fin | *hoxba* $^{+/-}$ ;*hoxbb* $^{+/-}$ | *hoxba+/-;b5b, b6b, b8b* gRNA-injected | -/- | +/- | frameshift (2bp ins) | 18bp del + 9bp insert | frameshift (5bp del) |
| 4 | No pectoral fin | *hoxba* $^{+/-}$ ;*hoxbb* $^{+/-}$ | *hoxba+/-;b5b, b6b, b8b* gRNA-injected | -/- | +/- | frameshift (5bp del) | 18bp del + 9bp insert | frameshift (2bp del) |
| 5 | No pectoral fin | *hoxba* $^{+/-}$ ;*hoxbb* $^{+/-}$ | *hoxba+/-;b5b, b6b, b8b* gRNA-injected | -/- | +/- | frameshift (5bp del) | frameshift(10bp del) | No mutation |
| 6 | No pectoral fin | *hoxba* $^{+/-}$ ;*hoxbb* $^{+/-}$ | *hoxba+/-;b5b, b6b, b8b* gRNA-injected | -/- | +/- | frameshift (5bp del) | frameshift(10bp del) | frameshift (1bp ins) |
| 7 | No pectoral fin | *hoxba* $^{+/-}$ ;*hoxbb* $^{+/-}$ | *hoxba+/-;b5b, b6b, b8b* gRNA-injected | -/- | +/- | frameshift (5bp del) | frameshift(10bp del) | frameshift (1bp del) |
| 8 | No pectoral fin | *hoxba* $^{+/-}$ ;*hoxbb* $^{+/-}$ | *hoxba+/-;b5b, b6b, b8b* gRNA-injected | -/- | +/- | frameshift (2bp ins) | frameshift (1bp insert) | frameshift (10bp del) |
| 9 | No pectoral fin | *hoxba* $^{+/-}$ ;*hoxbb* $^{+/-}$ | *hoxba+/-;b5b, b6b, b8b* gRNA-injected | -/- | +/- | frameshift (2bp ins) | 18bp del + 9bp ins | frameshift (5bp del) |
| 10 | No pectoral fin | *hoxba* $^{+/-}$ ;*hoxbb* $^{+/-}$ | *hoxba+/-;b5b, b6b, b8b* gRNA-injected | -/- | +/- | frameshift (5bp del) | frameshift(10bp del) | frameshift (1bp ins) |
| 11 | No pectoral fin | *hoxba* $^{+/-}$ ;*hoxbb* $^{+/-}$ | *hoxba+/-;b5b, b6b, b8b* gRNA-injected | -/- | +/- | frameshift (2bp ins) | 18bp del + 9bp ins | frameshift (5bp del) |
| 12 | No pectoral fin | *hoxba* $^{+/-}$ ;*hoxbb* $^{+/-}$ | *hoxba+/-;b5b, b6b, b8b* gRNA-injected | -/- | +/- | frameshift (5bp del) | frameshift(10bp del) | frameshift (1bp del) |
| 13 | No pectoral fin | *hoxba* $^{+/-}$ ;*hoxbb* $^{+/-}$ | *hoxba+/-;b5b, b6b, b8b* gRNA-injected | -/- | +/- | frameshift (2bp ins) | No mutation | frameshift (5bp del) |
| 14 | No pectoral fin | *hoxba* $^{+/-}$ ;*hoxbb* $^{+/-}$ | *hoxba+/-;b5b, b6b, b8b* gRNA-injected | -/- | +/- | frameshift (2bp ins) | No mutation | frameshift (10bp del) |
| 15 | No pectoral fin | *hoxba* $^{+/-}$ ;*hoxbb* $^{+/-}$ | *hoxba+/-;b5b, b6b, b8b* gRNA-injected | -/- | +/- | frameshift (5bp del) | frameshift(10bp del) | frameshift (1bp ins) |
| 16 | No pectoral fin | *hoxba* $^{+/-}$ ;*hoxbb* $^{+/-}$ | *hoxba+/-;b5b* gRNA-injected | -/- | +/- | frameshift (7bp del) | No mutation | No mutation |
| 17 | No pectoral fin | *hoxba* $^{+/-}$ ;*hoxbb* $^{+/-}$ | *hoxba+/-;b5b* gRNA-injected | -/- | +/- | frameshift (7bp del) | No mutation | No mutation |
| 18 | No pectoral fin | *hoxba* $^{+/-}$ ;*hoxbb* $^{+/-}$ | *hoxba+/-;b5b* gRNA-injected | -/- | +/- | frameshift (7bp del) | No mutation | No mutation |
| 19 | No pectoral fin | *hoxba* $^{+/-}$ ;*hoxbb* $^{+/-}$ | *hoxba+/-;b5b, b6b* gRNA-injected | -/- | +/- | frameshift (2bp del + 72bp ins) | frameshift (3bp del + 10bp ins) | No mutation |
| 20 | No pectoral fin | *hoxba* $^{+/-}$ ;*hoxbb* $^{+/-}$ | *hoxba+/-;b5b, b6b* gRNA-injected | -/- | +/- | frameshift (2bp del + 72bp ins) | frameshift (3bp del + 10bp ins) | No mutation |

**Table S2. The target-specific sequences of crRNAs used in this study**

| crRNA | Sequence | Reference |
|---|---|---|
| *hoxb4a* | 5'-GUUCACCCCCUGAAUAGUUC-3 | Maeno et al., Development (2024) |
| *hoxb5a* | 5'-UAUUUGGGGAGCUUGGCCAU-3' | Maeno et al., Development (2024) |
| *hoxb5b* | 5'-UCGACUACAACACCUAACGA-3' | Maeno et al., Development (2024) |
| *hoxb6b* | 5'-CGGGUAAACCGGGGUACAUG-3 | Maeno et al., Development (2024) |
| *hoxb8b* | 5'-UGACUGCCCAAGCUACACGC-3' | Yamada et al., Development (2021) |
| *hoxb4a* deletion | 5'-UUUUACCACUAUCUCAUCAA-3' | In this study |
| | 5'-AAAGUACGAGCAAGAUGUGU-3' | In this study |
| *hoxb5a* deletion | 5'-UAGCACCAAUGAUAAGAUAU-3' | In this study |
| | 5'-UAGAGCUUCACUUCAUACAA-3' | In this study |
| *hoxb5b* deletion | 5'-AAUUUACGACUGGUCAACAA-3' | In this study |
| | 5'-GACGUUUCAAUACAAGUUCA-3' | In this study |
| *hoxb4a-b5a* deletion | 5'-UAGCACCAAUGAUAAGAUAU-3' | In this study |
| | 5'-AAAGUACGAGCAAGAUGUGU-3' | In this study |
| *raldh2* | 5'-AAAAUAUUGACGACUCCCGG-3' | In this study |

**Table S3. The sequences of primers used for the genotyping in this study**

| Gene | Primer | Sequence | Reference |
|---|---|---|---|
| *hoxb4a* | *hoxb4a*-s | 5'-TCTAGCTTCTTCAATGTGTGGGGC-3' | Maeno et al., Development (2024) |
| | *hoxb4a*-as | 5'-TTCTTCTGCGGGTCAGATAGCG-3' | Maeno et al., Development (2024) |
| *hoxb5a* | *hoxb5a*-s | 5'-TCTCACAGAGCGAACAACTC-3' | Maeno et al., Development (2024) |
| | *hoxb5a*-as | 5'-GTAGTTTCCTCATCCAAGGG-3' | Maeno et al., Development (2024) |
| *hoxb5b* | *hoxb5b*-s | 5'-CAGATGAAACCAATGTCTCGTCCG-3' | Maeno et al., Development (2024) |
| | *hoxb5b*-as | 5'-TACCATGGCTAATGTGCAGCTTCC-3' | Maeno et al., Development (2024) |
| *hoxb6b* | *hoxb6b*-s | 5'-CTATTGGAAGTCTGGAAGACTCGC-3' | Maeno et al., Development (2024) |
| | *hoxb6b*-as | 5'-CGTTGACAGTCTCGTAGTGCTAAG-3' | Maeno et al., Development (2024) |
| *hoxb8b* | *hoxb8b*-s | 5'-GGGGAGTTGATCTTTCAAGCAATGA-3' | Yamada et al., Development (2021) |
| | *hoxb8b*-as | 5'-GTGGTAGAAATCCGGGAACTGA-3' | Yamada et al., Development (2021) |
| *hoxb4a* deletion | *hoxb4a* del-5'-s | 5'-CGTGACTTTCTATGGAGCTGGT-3' | In this study |
| | *hoxb4a* del-5'-as | 5'-CGATTCCCGGATAAGGAAATGG-3' | In this study |
| | *hoxb4a* del-3'-as | 5'-CGGGATAAAATGGGAACCAGAC-3' | In this study |
| *hoxb5a* deletion | *hoxb5a* del-5'-s | 5'-CGAAGCAGCAATTGGCACTTTGTC-3' | In this study |
| | *hoxb5a* del-3'-s | 5'-GAGAGTTAGCATGTGAGCTCTT-3' | In this study |
| | *hoxb5a* del-3'-as | 5'-CAGAGGACCATGAAACATAGGC-3' | In this study |
| *hoxb5b* deletion | *hoxb5b* del-5'-s | 5'-GGCACCATTGAGTTGAACCAAACG-3' | In this study |
| | *hoxb5b* del-3'-s | 5'-GTACAGTACAACTCGCACTACC-3' | In this study |
| | *hoxb5b* del-3'-as | 5'-CATTTCCTCGCGATGCAGGTATC-3 | In this study |
| *hoxb4a-b5a* deletion | *hoxb5a* del-5'-s | 5'-CGAAGCAGCAATTGGCACTTTGTC-3' | In this study |
| | *hoxb4a* del-3'-s | 5'-GTAGAACGTGGATTTTACCCGC-3' | In this study |
| | *hoxb4a* del-3'-as | 5'-CGGGATAAAATGGGAACCAGAC-3' | In this study |
